# Naturalistic language comprehension engages a cascade of widespread brain networks in the one second following comprehension

**DOI:** 10.1101/2024.12.17.629054

**Authors:** Min Kyung Hong, Andrew Janson, Prasanna Koirala, Tess Fotidzis, Sean Polyn, Katherine Aboud

**Author notes:** **Author Note:** This research was supported in part by a grant from NIH to KSA (DP5 OD031843). We would like to thank Ashley Jenkins, Adam Nassar, and Brooke Schnitzlein for help running the experiments. Corresponding author: Clair Min Kyung Hong.

## Abstract

Language comprehension (LC) is a cornerstone of human cognition, enabling the extraction of meaning from written and spoken communication with remarkable efficiency. Decades of neuroimaging research have identified the brain networks associated with LC, but limitations in spatial and temporal resolution have hindered a comprehensive characterization of the whole-brain (millimeter scale), real-time (millisecond precision) dynamics underlying this process. To overcome these constraints, we applied a fusion of multimodal brain imaging techniques (fMRI and EEG) in healthy adults (n = 30) to map the spatiotemporal progression of neural network engagement during LC in the one second following comprehension. Our findings reveal a cascade of brain network activations, beginning with the occipitotemporal perceptual word processing network (250 ms), followed by the temporoparietal semantic retrieval network (400 ms), the posterior default mode inferential network (500 ms), the frontotemporal semantic integration network (600 ms), and finally, a distributed goal-directed comprehension network (700 ms). Crucially, inferential processing emerged as a “hinge point,” linking early word processing to later higher-order networks. Efficient LC was associated with greater mediation by this inferential network and reduced reliance on top-down semantic integration. These findings provide evidence that naturalistic LC relies on rapid, dynamic interactions across widespread brain networks, with individual differences in LC reflecting specific subnetwork interactions. This work offers a framework for investigating the temporal evolution of distributed brain network dynamics in complex cognition across domains and clinical populations.

## Introduction

Naturalistic language comprehension (LC) is one of the most complex forms of human cognition, enabling us to extract meaning from written and spoken communication, form coherent representations of events, and navigate complex social interactions. This intricate process requires the rapid and flexible coordination of multiple brain networks, spanning perceptual, core language, and higher-order neurocognitive systems (Fedorenko et al., 2024; Church, 2023; Aboud et al., 2023; Bailey et al., 2018; Hagoort, 2013, 2017) Decades of neuroimaging research have revealed key regions and networks supporting LC, yet the dynamic interplay of these networks across space and time remains poorly understood, particularly during naturalistic language tasks. Understanding how distributed brain networks coordinate in real time is critical, as LC deficits are a hallmark of many developmental, neurological, and psychiatric conditions (Shaywitz et al., 2002; Petersen et al., 2013). However, existing methodologies have faced limitations: functional MRI (fMRI) excels in spatial localization but lacks temporal precision, while electroencephalography (EEG) captures rapid neural dynamics but offers limited spatial resolution. Magnetoencephalography (MEG) has bridged this gap in superficial cortical areas but struggles with deep brain structures and whole-brain networks. These constraints raise a fundamental question: how do brain networks dynamically interact in real time to support complex cognitive processes, such as efficient LC?

Previous research has shown that LC involves hierarchical processes from perceptual word recognition and semantic retrieval to the integration of meaning across sentences. These processes have been associated with distinct brain regions, including the occipitotemporal cortex for word-form perception, the middle temporal gyrus (MTG) for semantic retrieval, and the frontotemporal network for semantic control and integration. Event-related potential (ERP) studies have further identified characteristic temporal dynamics, such as the N400 (semantic processing; Kutas & Hillyard (1984); Kutas & Federmeier (2011); Lau et al. (2008)) and P600 (syntactic or reanalysis processing; Osterhout & Holcomb (1992); Brouwer et al. (2012); Friederici (2002); Hagoort et al. (1999)). Importantly, the degree of activation in these brain regions and temporal components corresponds with LC difficulties across clinical populations.

However, the spatiotemporal progression of these processes across whole-brain networks remains unclear. Perceptual and core language processes interact with the unfolding context and are driven by goal-directed cognition and working memory processes traced to nodes of the frontoparietal control network (FPN), as well as prediction and coherence-building processes linked to the default mode network (DMN). Numerous studies have associated DMN activation and connectivity with building coherent internal representations of ongoing events, including those represented in spoken or written discourse (Ferstl et al., 2008; Simony et al., 2016; Regev et al., 2013). P600, though traditionally associated with syntactic processing and controlled language processes, has also been linked to comprehension (Perfetti, 2007) and the engagement of higher-order cognitive networks, including the DMN and FPN (Aboud et al., 2023). The within- and between-network interactions of the DMN and FPN are specialized to content type and crucial for comprehension outcomes (Aboud et al., 2023). Despite the significance of higher-order networks in comprehending connected language, the temporal characteristics of the DMN and FPN subnetworks, and when they interact with perceptual and core language networks, remain poorly defined.

Notably, brain network dynamics across these processing levels are not modular: perceptual, core language, and higher-order brain networks dynamically interact (Fedorenko, 2014), and these interactions are facilitated by multifunctional “hub” regions (Aboud et al., 2023). Hubs are proposed to support efficient communication both within- and between-local brain networks, and consequently contribute to a wide range of functions (Friederici, 2011; Hagoort & Indefrey, 2014). However, the spatiotemporal characteristics of these hub areas have been largely determined through slow time-course examinations (Dronkers et al., 2017).

To address these gaps, we applied a novel fused fMRI-EEG analysis in healthy adults to map the spatiotemporal progression of brain network engagement during LC. This approach allowed us to resolve both where and when neural processes occur, providing unprecedented insight into the real-time, whole-brain dynamics of LC. By examining LC during naturalistic tasks (reading scientific texts) and comparing these to isolated word processing, we identified key network interactions that support efficient LC. Our findings offer a comprehensive framework for understanding how distributed brain networks coordinate to enable LC and lay the groundwork for applications across cognitive domains and clinical contexts.

## Methods

### Participants

Thirty-four right-handed adult participants were recruited from the community. All participants were native English speakers with normal or corrected-to-normal vision, no history of major psychiatric illness, and no contraindication to magnetic resonance imaging (MRI). Subjects came in for one visit at Vanderbilt University as part of an ongoing longitudinal study. During the visit, subjects participated in MRI, EEG, and paper/pencil testing sessions. To ensure that subjects had non-verbal IQ within the normal range and did not have dyslexia or oral language deficits, we administered the WAIS—matrices subtest (Drozdick et al., 2012), Woodcock Johnson Reading Mastery Test (WRMT) IV—Letter Word Identification (LWID) and Word Attack (WA) subtests (Schrank & Wendling, 2018), and the Peabody Picture Vocabulary Test (PPVT; Dunn (2019)). Behavioral metrics confirmed that subjects had typical IQ (minimum percentile score > 50; mean = 76.5 *±* 18.5), basic reading ability (minimum > 87 ss; mean = 104 *±* 8), and oral language ability (minimum percentile score > 17; mean = 75.6 *±* 16.3). Out of the original subject pool, *n* = 4 were excluded due to motion artifacts/excessive movement (*n* = 2), incidental brain abnormality finding (*n* = 1) and technical failure (*n* = 1). The final analysis included 30 adults (mean age = 25.59; 19 female). Participants gave written informed consent at the beginning of the study, with procedures carried out in accordance with Vanderbilt University’s Institutional Review Board (IRB). Participants received monetary compensation for behavioral and neuroimaging testing as per the study’s IRB.

### Stimuli

#### Passages

Five expository passages were developed based on the Soldier’s Manual and Trainer’s Guide. Using Coh-Metrix 3.0 (Graesser et al., 2014), the following measures were examined to ensure that all passages were matched in length and difficulty: word count (*M* = 9.75), syllable count (*M* = 1.49), word frequency (*M* = 2.02), word concreteness (*M* = 432), and Flesh Kincaid Grade Level (*M* = 5.7). To ensure equivalence of all measures across passages, measures for each of the five passages were individually compared to the bootstrapped mean of the remaining four passages. Passages were considered equivalent when measures were within a 90% confidence interval of the mean of the remaining passages.

### Text Features

From each passage, three component text features relevant to comprehension were examined at sentence level to interrogate the relationship between content-based cognitive demands and LC brain processes: (1) Verb Cohesion, (2) Deep Cohesion, and (3) Word Concreteness. Verb cohesion reflects the degree to which there are overlapping verbs within the text. When there are repeated verbs, the text includes enhanced cross-sentence semantic and structural priming. Deep Cohesion measures the extent to which a reader must infer meaning in the text. If the text includes intentional and causal connectives when there are logical and causal relationships within the text, these connectives enable the reader to form a more coherent and deeper understanding of the causal actions, processes, and events in the text. When a text contains many relationships but does not contain those connectives, the reader must infer the relationships between the ideas within the text. Lastly, word concreteness measures the imageability of the content and is related to the ease of building an internal representation of the text (Sabsevitz et al., 2005; Binder et al., 2005). Texts with more concrete and imageable content words are easier for readers to process and comprehend. For example, “The forest was lush and verdant” is more concrete than “The concept was abstract and complex”. Abstract words typify concepts that are difficult to visualize. Therefore, texts that contain more abstract words are more challenging to understand.

#### Words

The word baseline of a passage consisted of scrambled words from the other four passages (i.e., Passage A word baseline consisted of scrambled words randomly sampled from Passages B, C, D, and E). Stimuli were presented one word at a time, and words were matched in length and presentation time to the passage paragraphs. The word baseline was inspected to ensure that the scrambled words did not form a sentence.

### Experimental Design & Procedures

Subjects read Passages and isolated Words while in the MRI, and in a separate session, during the EEG. Stimuli selection was randomized so that each subject viewed two randomly selected Passages (out of 5) and associated Word stimuli. For each subject, stimuli was matched across the MRI and EEG sessions. Passages and Words were comprised of four paragraphs each in a single run (five sentences or 50 words per paragraph), and conditions were ordered as follows: Words paragraphs 1 & 2, Passage paragraphs 1 & 2, Words paragraph 3 & 4, Passage paragraphs 3 & 4. During each session, Passages and Words were presented one word at a time (fig. 1). Stimuli appeared in white letters on a black background, using Comic Sans MS font, size 32. Based on previous literature on reading speed (see Brysbaert (2019) for a review), each word was displayed for 500 ms - 600 ms, varying based on the number of syllables in the word to provide adequate time for encoding and processing of text. A single-syllable word was presented for 500 ms, and an additional 25 ms was added for each subsequent syllable. For example, the word “hot” was presented 500 ms, while the word “inhalation” was presented for 575 ms. Each word presentation was followed by a 100 ms pause between words. During the Passage condition (B. Passage Paragraphs), the sentence’s terminal word was followed by a 1000 ms break (indicated by a plus sign). To monitor whether participants attended to all stimuli, 5% of the stimuli within each task block were randomly repeated on two consecutive screens. Participants pressed a button with their right thumb when they detected a phrase repetition or a symbol configuration repetition. All participants had high accuracy (*>* 70%) for the repetition flag, confirming that they stayed on task.

**Figure 1.**
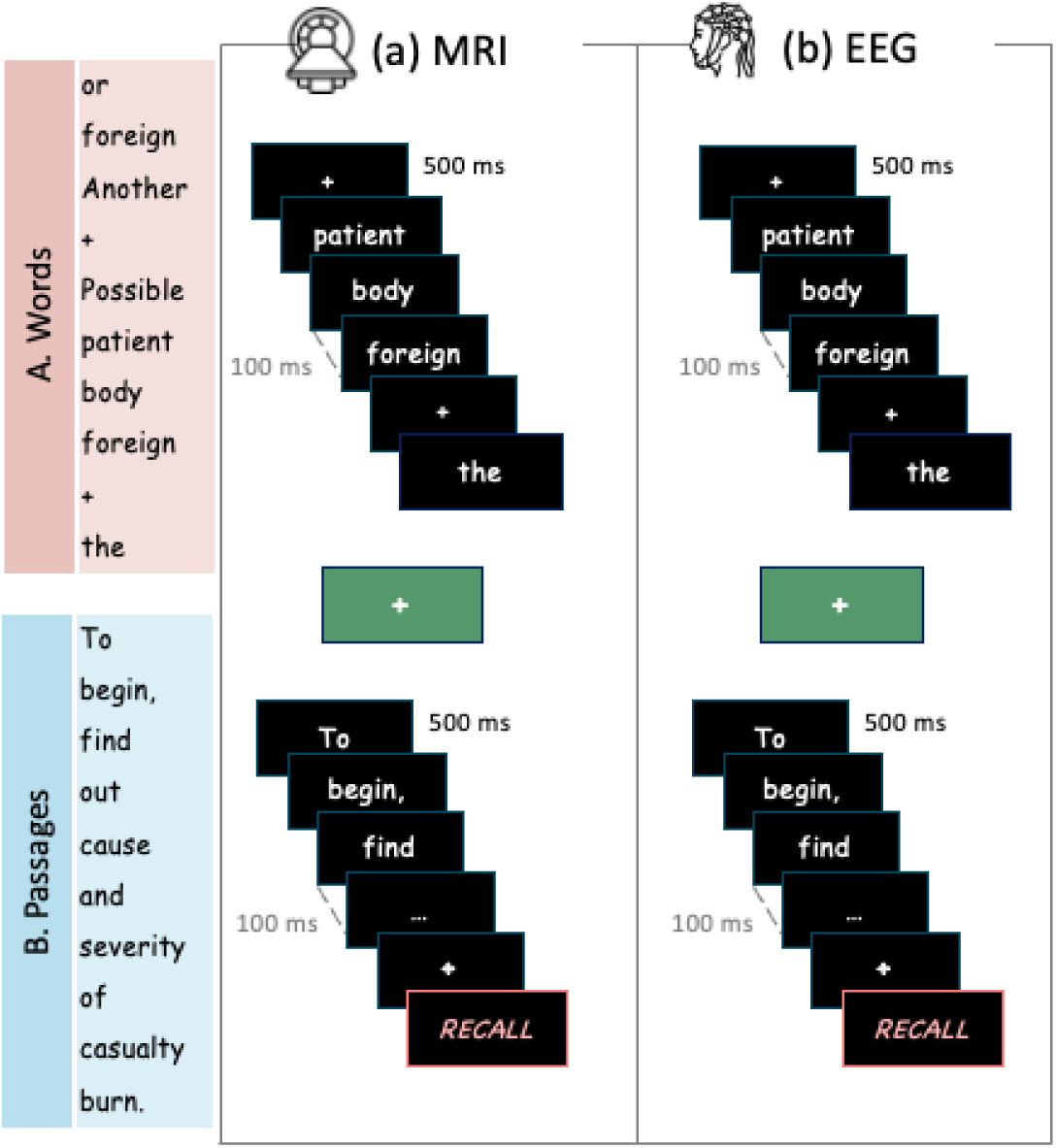
[Experimental design overview. (a) MRI: In the scanner, participants viewed alternating blocks of scrambled words and passage paragraphs. Each screen was displayed for 500+ ms with 100 ms inter-stimulus intervals. A single run sequence consisted of: (1) scrambled words, (2) the first two paragraphs from a passage, (3) a new set of scrambled words, and (4) last two paragraphs of the same passage. The word count in scrambled blocks matched subsequent passage paragraph blocks. (B) EEG session: Immediately after MRI, the same stimuli were presented in the same order in a separate EEG setting.

### fMRI Data Acquisition, Preprocessing and First-Level Analyses

Imaging was performed on a Philips Achieva 3T MR scanner with a 32-channel head coil. Functional images were acquired using a gradient echo planar imaging sequence with 40 (3 mm thick) slices with no gap, consisting of 2 runs, each approximately 6 minutes (255 dynamics per run). Additional imaging parameters for functional images included: TE = 35 msec (for optimal BOLD contrast at 3T), FOV = 240 x 240 x 120 mm, slice thickness = 3 mm with 0 mm gaps, 75 degree flip angle, TR = 1300 msec, and a matrix size of 80×80 (interpolated), yielding 3 mm³ isotropic voxels. All functional data were analyzed using MATLAB R2022b and SPM12 (Friston et al., 1994). The functional data for each participant were slice-timing corrected, aligned to the mean functional image, co-registered to the subject’s T1 structural image, normalized to MNI space, and spatially smoothed with a 6 mm FWHM Gaussian filter. In our first-level analysis, standard regression models were created using an estimated HRF for each condition. The six motion parameters (3 translations and 3 rotations) were included as nuissance regressors in the design matrix. Outlying volumes, as determined by ART (Whitfield-Gabrieli; http://www.nitrc.org/projects/artifact_detect/), were also included in the design matrix as regressors of no interest. All 30 subjects in our analysis showed less than 10% average motion outliers and less than 15% motion outliers in any individual run.

### EEG Data Acquisition and Preprocessing

All EEG data were acquired at the Vanderbilt Kennedy Center, using a 128-channel geodesic sensor net (EGI, Inc., Eugene, OR). Data were sampled at 250 Hz with filters set to 0.1–30 Hz. The vertex was used as the reference during data acquisition. Data processing was completed using NetStation and MATLAB (EEGLAB toolbox). EEG data were segmented into epochs of 1000 ms, starting 100 ms before the onset of the critical word. For all conditions, the critical word was in the sentence-final position. Recordings were re-referenced to an average reference. Automated artifact rejection and ICA labeling were used to remove ocular and muscle artifacts, including eye movements and trials that were identified through automated and manual artifact identification processes. Contaminated electrodes in each trial were rejected, and trials with more than ten rejected electrodes were excluded from analysis (n = 1 subject was excluded due to excessive motion artifacts). Within each participant, pre-processed time signals for each sentence were averaged into paragraphs. These paragraph averages were input into the joint ICA pipeline (Electrodes: 7, 106, 31, 55, 80, 54, 62, 79).

### fMRI – ERP joint Independent Component Analysis

Joint ICA with fMRI and EEG is a fused multimodal analytical approach that identifies co-occuring patterns across brain imaging modalities. By taking fMRI spatial signals and EEG temporal signals, this simultaneous decomposition allows us to reliably estimate temporal signals in the 1 second following comprehension and their corresponding spatial patterns, revealing how the rapid electrical responses (ERPs) map onto more slowly changing metabolic activity in specific brain regions (fMRI). Following protocols used in Aboud et al. (2023), we used the Fusion ICA Toolbox (FIT) in MATLAB to conduct fMRI and EEG joint ICA (see also Calhoun et al., 2006; Mijović et al, 2012). As shown in Figure A1, the input for jICA is a subject *×* data input matrix where each subject’s spatial fMRI map and the ERP component time course are concatenated into a single matrix. The fMRI and ERP input data are first-level contrast maps and grand average time courses (averaged across centroparietal electrodes).

Fusion analysis was conducted using the Fusion ICA Toolbox (FIT) in MATLAB, following protocols established by Calhoun et al. (2006) and Mijović et al. (2012). These protocols were designed for parallel fMRI/EEG acquisition, which is considered more suitable for this approach than simultaneous acquisition (Calhoun et al., 2006; Mijović et al., 2012). In jICA, independent components for fMRI and EEG are estimated simultaneously. Unlike other multimodal analysis methods, jICA allows the spatial and temporal components of EEG and fMRI to influence each other, making it a true *fused* data analysis approach. Spatial fMRI maps and ERP component timecourses for each paragraph and condition were concatenated into a subject-by-data input matrix. For example, the fMRI and ERP data for one input would be the first-level contrast map and grand average timecourse for a subject’s Passage Paragraph 1. Thus, participants had up to 8 spatiotemporal inputs (four for word baseline, four for passage). In our dataset of n = 30, 90% of participants had seven or more total input data per run. ERP timecourses were upsampled using cubic spline interpolation to match the dimensionality of the spatial fMRI vector. The model assumes that ERP peaks and BOLD responses change similarly across subjects, providing robust, high-quality data decompositions validated across various cognitive substrates and populations. The jICA algorithm produces group-level, joint independent components that include information from both modalities (i.e., each component has both an ERP timecourse and a spatial map). Condition-specific maps and timecourses are back-reconstructed to identify how each condition contributes to the cross-condition components. The strength of this contribution is indicated by a subject- and condition-specific scalar parameter loading, which measures the component signal’s strength within each subject and condition and can be used to statistically identify differences between conditions (see statistical approach). For all jICA results, spatial maps are thresholded at z > 2.5 (equivalent to p < 0.005). Multiple comparison correction of the ICA decomposition was not necessary as it is a multivariate, not univariate, approach.

### Joint ICA Parameters

The Infomax algorithm was employed to identify joint components in our MRI-EEG data fusion analysis (Bell & Sejnowski, 1995; Calhoun et al., 2006). Internal bootstrapping was run via ICASSO (n = 50 iterations) to determine the stability of each component. All components had a stability Iq index above 0.95 (mean = 0.97).

### Component number calculation

To determine the optimal number of components, we followed protocols established in (Aboud et al., 2023; Himberg et al., 2004; Turner et al., 2012; James et al., 2014). First, we ran joint ICA in 15 components, which we have previously found to be a stable and replicable component number in language processing (Aboud et al., 2023). From this, Principal Component Analysis was run for the combined fMRI and EEG data to determine the amount of cumulative variance explained by the number of components. We found that 95 percent of the cumulative variance was explained by 13 components (as done in Calhoun et al. (2009); Mijović et al. (2010)). ICASSO bootstrapping was used to determine component stability. All components demonstrated a stability index (Iq) > .95, indicating that the components were robust to randomized start points in the ICA algorithm. To validate our selection, we conducted a qualitative comparison of our primary results with adjacent component numbers (e.g., *k* = 11, 13, and 15) and found that they yielded highly the same ERP peak subdivisions in the N400 and P600 range. Additionally, we explored the effects of extreme component numbers.

Low component numbers (e.g., *k* = 8) resulted in the merging of significant subcomponents. For instance, the P600 component appeared as a single component with multiple peaks around 500 ms - 700 ms. High component numbers (e.g., *k* = 50), on the other hand, led to an excessive division of relevant subcomponents. This combined approach identified the point at which additional components contributed minimally to the total variance explained while retaining the most significant sources of variation in our data and avoiding over-fitting.

### Validation Analysis

To validate the robustness of our findings, we conducted a permutation analysis with a leave-out cross-validation approach. Specifically, we performed 10 iterations of 80/20 splits of the subject data, where 20 percent of the subjects were randomly left out in each iteration. For the remaining 80 percent of subjects, we reran the full analysis. We manually identified the 5 temporal components that aligned with JCs 1-5, then generated back-reconstructed fMRI images for Passages and Words across each of the five identified components. These back-reconstructed images were thresholded at *Z >* 1.5 and masked by the original joint component spatial map. To quantify the consistency of these findings, we calculated the intraclass correlation coefficient (ICC) (McGraw & Wong, 1996) across the outputs of the 10 permutations. ICC values were computed for each component by comparing the fMRI images masked by the original fMRI findings (thresholded at z > 1.5). For Passages, the ICC values were as follows: JC1: ICC = 0.743 (*p <* 0.001), JC2: ICC = 0.807 (*p <* 0.001), JC3: ICC = 0.772 (*p <* 0.001), JC4: ICC = 0.810 (*p <* 0.001), JC5: ICC = 0.740 (*p <* 0.001). For Words, the ICC values were only moderate or strong in JC1 and JC2, with JC1: ICC = 0.839 (*p <* 0.001), JC2: ICC = 0.763 (*p <* 0.001), JC3: ICC = 0.033 (*p* = 0.0042), JC4: ICC = −0.143 (n.s.), JC5: ICC = 0.566 (*p <* 0.001). These results demonstrate a high degree of consistency across permutations for Passages, and consistency across only early components (JCs 1 and 2) for Words.

### Component Inclusion

For each of the 13 components, each ERP had to meet signal criteria to ensure that the ERP timecourse did not represent noise. Specifically, through a manual inspection and using findpeaks MATLAB function, components were excluded from further analysis if they showed excessive peaks or no clear peak. Additionally, back-reconstructed components were only considered if they met validation criteria of ICC > 0.7 (p < 0.001).

### Statistical Approach

In order to examine how jICA components mapped onto condition and text features, paragraph-level Joint ICA component weights from each subject were examined within a linear mixed-effects regression (LMER) framework in R (lme4 package, (Bates et al., 2015)). This approach provided methodological flexibility to account for all available information (e.g., within-subject trial-to-trial variability in brain signals in response to varying text features, varying number of inputs per subject, and variability in reading abilities across subjects).

First, we conducted a series of linear mixed-effects regression analyses on jICA component weights, examining the main effect of condition (Words vs. Passage), with subject vocabulary ability (PPVT standardized score) as a covariate, and their interaction to control for individual differences that may modulate the effect of condition, with a random intercept for subject (e.g., *JointComponent_i_* = *β*_0_ + *β*_1_Condition + *β*_2_PPVT + *β*_3_(Condition *×* PPVT) + (1*|Sub ject*) + *ε*).

Second, to examine which joint components’ weights were predictive of text difficulty, we conducted additional analyses that include the five joint component weights during passage reading as independent variables and the text feature as a dependent variable (subjects as random effects) (e.g., *VerbCohesion* = *β*_1_JC1 + *β*_2_JC2 + *β*_3_JC3 + *β*_4_JC4 + *β*_5_JC5 + (1*|Sub ject*) + *ε*).

Third, to examine comprehension efficiency, the Gates MacGinite Reading Comprehension subtest was administered, in which participants read short texts (1–2 paragraphs in length) and answered questions about the story: the test consisted of 11 stories with a total of 48 questions, which participants completed within a 35-minute time limit. To test how engagement of the five joint component networks was modulated by comprehension efficiency, each participant’s passage jICA weight (dv) was examined with Gates MacGinite standardized reading scores as an independent variable with random intercept for subject. (e.g., *JointComponent_i_* = *β*_0_ + *β*_1_GM + (1*|Sub ject*) + *ε*).

Lastly, to better understand the dynamics of how these five joint components interact to give rise to LC, we conducted an exploratory factor analysis with Passage jICA weights using the minimum residual extraction method with oblimin rotation. First, we conducted preliminary statistical assessments to determine if EFA was appropriate. The Kaiser-Meyer-Olkin (KMO) measure confirmed sampling adequacy for the analysis with an overall metric of 0.74 and individual KMO values ranging from 0.72 to 0.78. Concurrently, Bartlett’s test of sphericity, *χ*^2^ = 258.03, *p <* 0.001, *d f* = 10, indicated that correlations between items were sufficiently large for a factor analysis. Based on the three factors obtained frmo EFA, we conducted a GLM mediation analysis with Factor 1 as a covariate, Factor 2 as DV, and Factor 3 as the mediator.

## Results

We examined 30 subjects (mean age 25.6 years, 11 male) as they read passages and isolated words while in the MRI and, in a separate session, during EEG. Passages were matched on difficulty and presented one word at a time; Words were matched in duration as Passages and consisted of randomly scrambled passages. To ensure attention, 5 percent of words in both conditions were repeated on adjacent screens, and subjects were instructed to press a button upon seeing a repeated word. Subject-level fMRI spatial maps and ERP time courses (averaged across trials) were generated and then input into joint independent component analysis (jICA). JICA is a fused brain imaging approach that allows for co-analysis of fMRI hemodynamic signals and EEG electrophysiological signals to reliably estimate the temporal progression of brain network activations at the millisecond and millimeter scale. In this approach, ERP component peaks can be used to order fMRI spatial maps in the one second following LC. Joint ICA produced 13 spatiotemporal components.

### Spatial and temporal signatures of naturalistic language processing at 100ms interval

First, we identified components specific to naturalistic LC. A linear mixed-effects model was run comparing Passage and Word component weights, while controlling for subject word reading ability. This analysis identified five components that were significantly greater in Passages versus Words, and which met filtering criteria (see Methods). The main effects of condition were as follows (Passages > Words, see tables B1 and C1 for full list of results): Joint Component 1 (β = 0.09, SE = 0.03, *t* = 2.90, *p* = 0.004), Joint Component 2 (β = 0.07, SE = 0.03, *t* = 2.14, *p* = 0.032), Joint Component 3 (β = 0.07, SE = 0.04, *t* = 1.97, *p* = 0.049), Joint Component 4 (β = 0.10, SE = 0.04, *t* = 2.66, *p* = 0.008), Joint Component 5 (β = 0.10, SE = 0.05, *t* = *−*2.27, *p* = 0.023). As shown in Figure 2, the five components fell within the canonical time window of LC (200-800 ms after the critical word). To identify which cognitive processes modulated these components, we leveraged text feature metrics in Passages, and ran linear mixed-effects models to identify which joint components’ weights were predictive of text feature difficulty (see Methods). Text features included three canonical component text measures found to capture the ease of LC subprocesses in connected texts (see Figure 2f): Verb Cohesion (linguistic processing demands of text), Deep Cohesion (conceptual inferential demands in text), and Word Concreteness (visualization demands of text).

**Figure 2.**
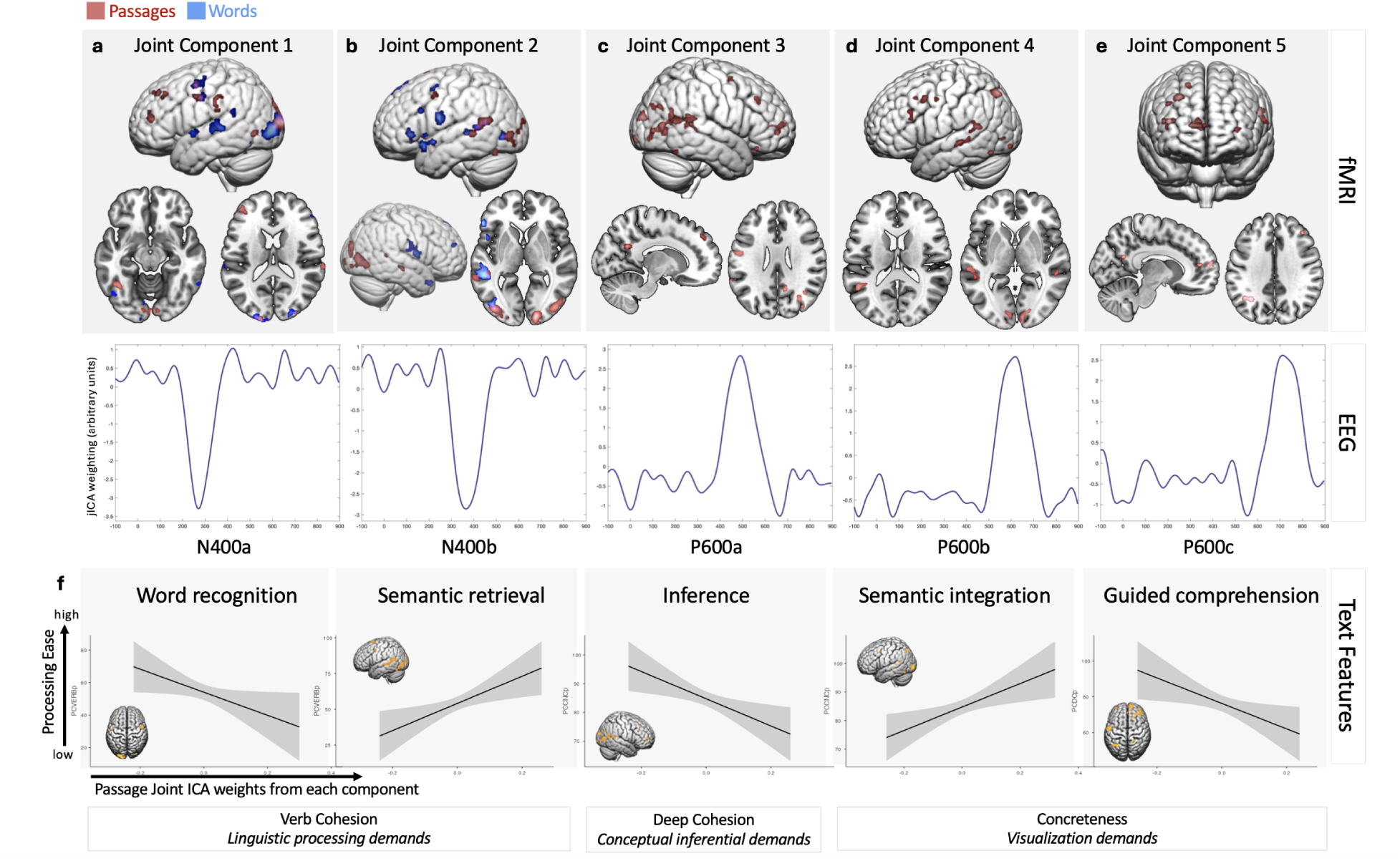
Five spatio-temporal joint ICA components (Red = Passages, Blue = Words, images displayed at z > 2.5). Spatial maps of Words & Passages in the N40O time window: (a) Joint Comp. 1, word recognition network (250 ms, L fusiform & STG);(b) Joint Comp. 2, semantic retrieval network (400 ms, MTG); Spatial maps of Passages from P600 time window: (c) Joint Comp. 3, inference network (500 ms, posterior DMN); (d) Joint Comp 4, semantic integration network (600 ms, L posterior MTG, L IFG; (e) Joint Comp. 5, guided comprehension network (700 ms; L AG, MFG, Precuneus) (f) Modulation of each joint component engagement based on text feature demands during passage reading. Significant relationships were observed in the following: JC1 & JC2 with verb cohesion (linguistic demands), JC3 with deep cohesion (conceptual inference demands), JC4 & JC5 with concreteness (visualization demands).

#### Joint Component 1: Word recognition

With ERP peak at around 250ms, JC1 for Passages was localized to a distributed network of frontal, temporal, and primary sensory areas linked to perceptual word processing (Fig. 2a): left fusiform gyrus (BA 37), bilateral IFG (BA 45), left middle frontal gyrus (BA46), right STG (BA41), and bilateral middle occipital regions (BA 19). A similar, but more focused, pattern was observed for Words: temporal (left MTG/STG, BA21 and BA22) regions, as well as occipital areas (bilateral middle/superior occipital, BA19). During passage reading, JC1 showed a significant negative relationship with verb cohesion (β = *−*0.71, *SE* = 0.35, *t* = −2.06, *p* = 0.041). The negative relationship suggests that greater engagement of JC1 word recognition processes was associated with low verb cohesion, potentially reflecting increased lexical access and integration demands.

#### Joint Component 2: Semantic retrieval

With an ERP peak around 400ms, JC2 for Passages was localized to canonical word meaning areas, including the left middle temporal gyrus and left temporoparietal junction (TPJ), along with primary visual areas acc(Fig. 2b). Words showed overlapping activation of the left middle temporal gryus, as well as a more widespread pattern in the language network, engaging temporal, frontal, parietal, and visual areas. This included bilateral MTG (BA 21, predominantly on the left side), left STG(BA 22), bilateral temporal pole (BA38), bilateral IFG (BAs44,45), left middle and superior medial frontal regions, left fusiform gyrus (BA 37), and left middle and superior occipital areas. JC2 showed a significant positive relationship with verb cohesion during passage reading (*β* = 0.95, *SE* = 0.35, *t* = 2.69, *p* = 0.008): greater engagement of the semantic retrieval processes reflected by JC2 was associated with higher verb cohesion, indicating more efficient lexical access and semantic integration for more cohesive text.

#### Joint Component 3: Inference

With a positive peak around 500ms, JC3 marked a shift into higher order brain networks during Passages, but Words did not robustly map on to the component (Fig. 2c). For Passages, JC3 mapped onto right-dominant posterior DMN, including precuneus, PCC, bilateral temporoparietal junction/right angular gyrus, middle frontal and inferior orbitofrontal gyri, and STG. JC3 during passage reading significantly predicted variance in text deep cohesion (β = *−*0.71, *SE* = 0.30, *t* = −2.34, *p* = 0.020). This negative relationship indicates that with higher conceptual inferential demands in text, characterized by lower deep cohesion, led to greater engagement of the syntactic-semantic reappraisal processes in JC3.

#### Joint Component 4: Semantic integration

With a positive ERP peak around 600ms, JC4 for Passages localized to the canonical frontotemporal semantic integration network, while Words did not robustly map on to the component. Passage activations included left posterior MTG (BA 21), left IFG/dlPFC, and bilateral superior parietal lobule (BA 7) (Fig. 2d). The left middle temporal gyrus has been implicated in semantic processing and integration, while the superior parietal lobule has been associated with attentional and executive control processes involved in language comprehension. During Passages,JC4 significantly predicted variance in word concreteness (β = *−*0.47, *SE* = 0.17, *t* = −2.70, *p* = 0.007). Word concreteness reflects the meaningfulness and imageability of content words, which can influence the visualization demands placed on the reader. The negative relationship indicates that more abstract sentences (lower in concreteness and thus more difficult to process) elicited greater engagement of JC4. This finding is consistent with previous work on the P600 (Kuperberg, 2007; Shen et al., 2016) as a signal associated with semantic and syntactic integration processes.

#### Joint Component 5: Conceptual coherence/guided comprehension

The final joint component for Passages had a positive peak at 700 ms, and mapped on to key nodes in two higher-order networks: the DMN and the FPN. In particular, the component showed midline DMN (mPFC, PCC, and PCU), the left dorsal AG/IPS, and coactivation in the medial and lateral prefrontal cortex (Fig. 2e). In addition, Passages showed activations in the bilateral STG, insula, and sensory motor areas. Words did not robustly map on to this component. Temporally, this component had a late positive peak latency around 700 ms post-stimulus, suggesting its involvement in higher-level conceptual integration and coherence processes during guided language comprehension. JC5 during passage reading showed a significant positive relationship with word concreteness (β = 0.41, *SE* = 0.15, *t* = 2.72, *p* = 0.007). This indicates that this component was more engaged for more concrete and imageable text are easier to integrate into a coherent mental representation.

#### Component hubs

Next, we identified component hubs through examining overlapping areas across components (Fig 3). We used boolean conjunctions across components. We found three main hubs in the reading, language, and higher order areas: left MTG (*JC*2 *∩JC*4; BA 21/22); left TPJ (*JC*2 *∩JC*3 and *JC*2 *∩JC*4;), and right PCC with *JC*3 *∩JC*5; BA31). These findings demonstrate that previously identified key areas in the perceptual word reading, core language and default mode network are interleaved across the timecourse of LC-related brain activations. Each hub area co-occurs with nodes from other joint networks linked to diverse cognitive demands, suggesting multifunctional roles in these densely connected areas.

**Figure 3.**
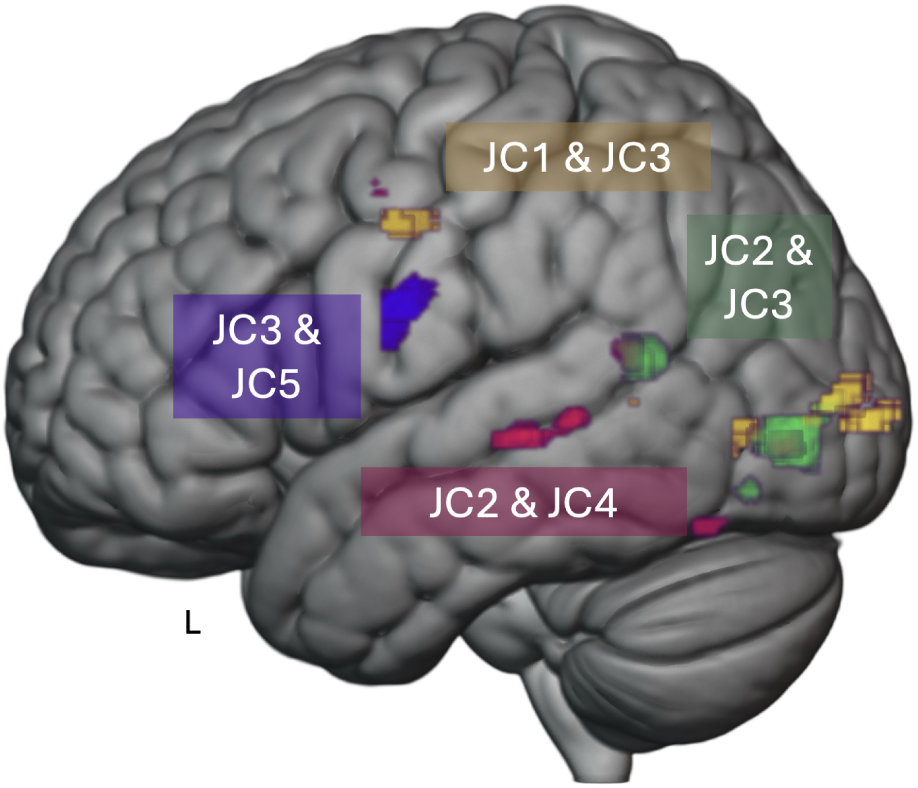
Joint Component Hubs: Overlapping areas across the five joint components were identified to examine how joint components from different timepoints may show up on the same area, revealing intersections of brain regions that potentially support multifunctional cognitive processing across network dynamics. The main overlapping areas were (1) JC2 ∩JC4 (red): MTG (BA 21/22) and TPJ, (2) JC2 ∩JC3 (green): TPJ, (3) JC3 ∩JC5 (blue), right PCC and primary sensory areas, and (4) JC1 ∩JC3 (yellow), primary sensory and visual areas.

#### Relationship Across Networks: JC3 as a bridge between lower- and higher-order LC processes

We were next interested in how joint components related to one another, i.e. do certain components cluster together. To examine this, we performed an exploratory factor analysis, which identified three latent factors corresponding with early, middle, and late temporal signals. Underlying the five JCs (see Methods for details), we found: Factor 1 with JC1, JC2, JC3, Factor 2 with JC4 and JC5, and Factor 3 with JC3. This three-factor structure highlights the shared yet distinct roles of each component. Early perceptual and core word processing from the two early N400 components are captured in Factor 1 (JC1 & JC2), the transition from local word-level processing to more global, context-level processing is reflected in JC3 as Factor 1 and Factor 3, and LC-specific executive processes and higher-order comprehension from the two late P600 components (JC4 & JC5) are captured through Factor 2. Interestingly, these factors also converged with our text feature findings where JC1 & JC2 were jointly predictive of linguistic processing demands, JC4 & JC5 sharing visualization demands and JC3 uniquely predicting conceptual inference demands. Based on these findings, we hypothesized that JC3 may be the bridge between early word-level semantic processes and later global comprehension processes. A mediation analysis with the JC1 & JC2 as a covarite, JC3 as a mediator, and the JC4 & JC5 as dependent variable revealed a significant partial mediation: we observed significant direct (β = 0.35, [*z* = 5.41, *p* < 0.001), indirect effect (β = 0.11, *z* = 3.37, *p* < 0.001), and total effects (β = 0.46, *z* = 7.64, *p* < 0.001).

### What does expert naturalistic language processing look like?

Next, we tested how engagement of the five joint component networks was modulated by expertise, i.e., we were interested in determining which spatiotemporal dynamics promote more *efficient* LC processes. First, we examined how LC ability corresponded with (1) component weights, and (2) the relationship between adjacent components (e.g. JC1 weights - JC2 weights). We found a significant negative relationship between participants’ reading ability (defined by standard scores from Gates MacGinite passage comprehension subset) and JC4, a top-down semantic integration network (β = *−*0.56, *SE* = 0.21, *t* = −2.69, *p* = 0.011) (Fig. 4a). We next asked whether there may be a trade off between the bottom-up DMN (JC3) and top-down semantic control (JC4) component engagement during passage reading in more efficient comprehension. Specifically, the ability to rely more on the bottom-up JC3 component compared to the top-down JC4 component may be indicative of more automatic rather than effortful processing (Derawi et al., 2022). Consistent with this idea, we found that participants’ reading ability significantly predicted the difference between JC3 and JC4 (*β* = 0.619, *SE* = 0.23, *t* = 2.68, *p* = 0.012)(Fig. 4b): better readers exhibited a larger difference, with higher JC3 and lower JC4.

**Figure 4.**
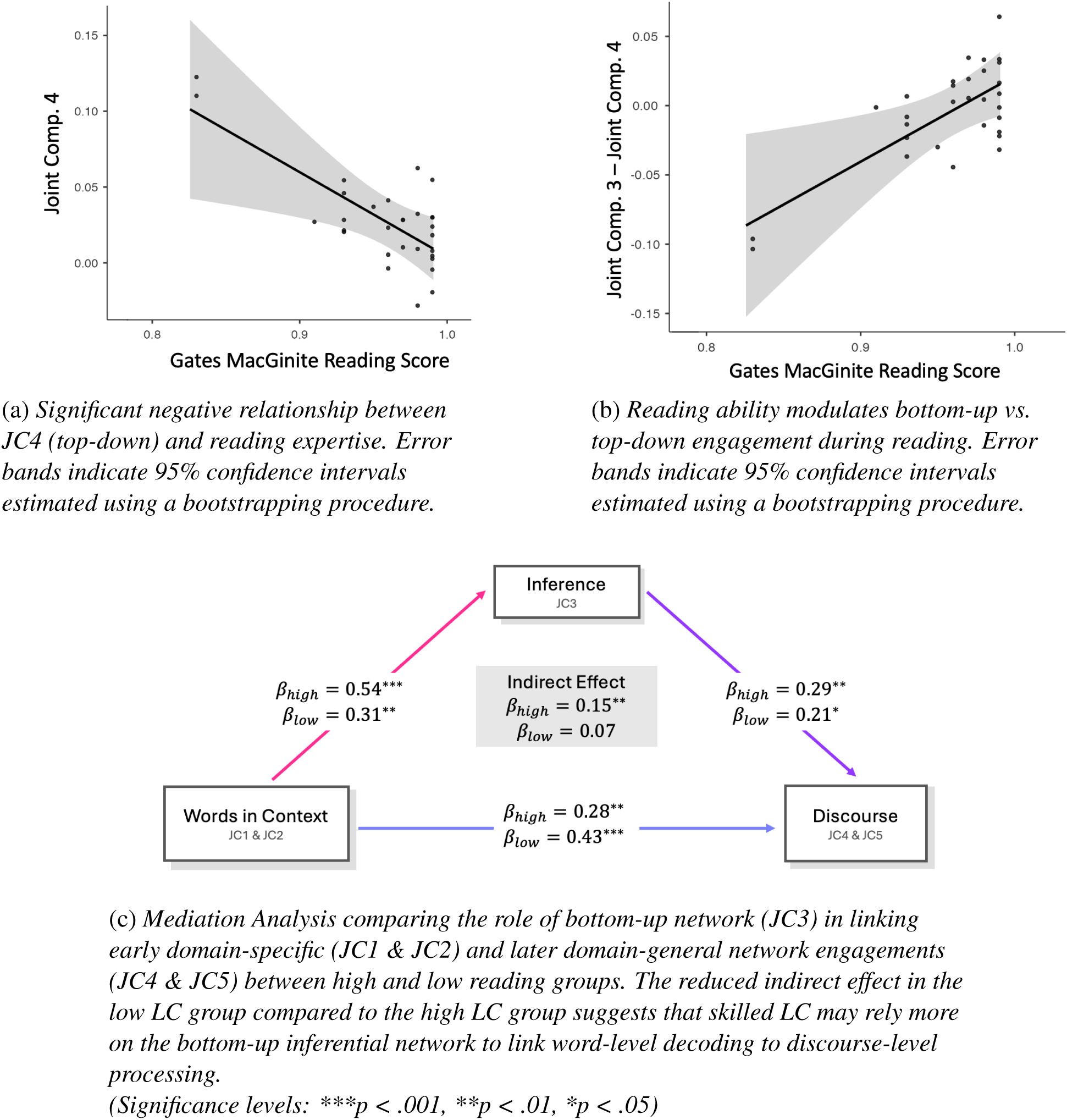
Expertise and top-down vs. bottom up engagement.

Based on the factor analysis results and the roles of each joint component, we next asked whether the general JC3 mediation effects described above were driven by “expert” readers, hypothesizing that the bottom-up DMN mediation processes would promote more efficient processing and consequently be a marker for higher LC ability. To test our hypothesis, we split participants into two groups, low and high, based on their reading comprehension scores (median split). We then conducted a mediation analysis on each group, assessing the extent to which JC3, the bottom-up network, accounted for the link between the early domain-specific (JC1 & JC2) and later domain-general network engagements (JC4 & JC5). While both groups showed significant direct (high: β = 0.28, *z* = 3.01, *p* < 0.001 vs. low: β = 0.43, *z* = 4.68, *p* < .001) and total effects (high: β = 0.43, *z* = 5.35, *p* < 0.001 vs. low: β = 0.49, *z* = 5.05, *p* < .001), the indirect effect was reduced in low group compared to the high group (high: β = 0.15, *z* = 2.86, *p* = 0.004 vs. low: *β* = 0.07, *z* = 1.87, *p* = 0.06). The reduced indirect effect in the low reading group suggests that skilled LC may be marked by reliance on the bottom-up inferential network to link word-level decoding to discourse-level processing.

## Discussion

Naturalistic LC is fundamental to human cognition. The current study provides a comprehensive framework for understanding the spatiotemporal dynamics of LC through fused fMRI and EEG. By addressing long-standing spatial and temporal limitations in neuroimaging, we reveal a cascade of neural network exchanges that span perceptual, core language, and higher-order cognitive processes. These findings advance our understanding of how distributed brain networks dynamically interact in real time to support complex cognition.

### Dynamic Interactions Between Brain Networks During LC

We found that naturalistic LC is marked by rapid interactions between perceptual, core language, and higher-order cognitive brain networks. We identified five LC brain networks with unique spatial, temporal, and linguistic characteristics: (1) early activations of perceptual word recognition areas at 250 ms (JC1); (2) temporal contextual meaning retrieval areas at 400 ms (JC2); (3) posterior DMN connectivity at 500 ms (JC3); (4) a frontotemporal language network at 600 ms (JC4); and (5) cross-network connectivity between core DMN and FPN nodes at 700 ms (JC5). These temporal signals corresponded to canonical EEG components, including the N400 and the P600, with overlapping and distinct localizations for Passages versus isolated Words. This suggests that these temporal signals are comprised of subsignals with independent localizations and that their activation distributions vary by task demands.

The first two components corresponded to the N400, an EEG signal related to semantic cognition (Kutas & Federmeier, 2011). JC1 mapped onto a perceptual word identification network, including in the left occipitotemporal cortex related to rapid visual word identification (Dehaene et al., 2005; Vinckier et al., 2007). Additional activation for Passages emerged in the SMG (phonological word processing) and lateral PFC (articulatory recoding and attentional control)(Pattamadilok et al., 2010; Bitan et al., 2005). Isolated Words elicited bilateral STG activation, reflecting phonological processing without context (Hickok & Poeppel, 2007). These findings suggest that JC1 supports dynamic perceptual word identification processes influenced by contextual linguistic features. JC2, active at 300 ms, localized to posterior temporal and temporoparietal hubs in Passages related to semantic retrieval (Seghier, 2013, 2022; Jefferies, 2013), while isolated Words elicited broader activations extending into frontal regions, indicative of more effortful binding of word meanings without context (Hagoort, 2005). Higher verb cohesion in texts increased engagement of JC2 hubs, demonstrating that predictive contextual processing in Passages may allow for greater spreading activations in the semantic retrieval network.

Following early perceptual and core word processes, the third joint component (JC3) was the first in the cascade to fall within the P600 window, an EEG signal related to syntax, semantic control, and comprehension (Laszlo & Federmeier, 2011; Brouwer et al., 2012). Interestingly, JC3 showed overlapping activation in the left TPJ from JC2 but now coupled with core nodes of the posterior DMN, including the PCC and right TPJ. Consistent with prior literature on the right TPJ and DMN, this component was significantly related to the inferential demands of the text (Seghier, 2013; Saxe et al., 2006). We additionally found that JC3 significantly partially mediated the relationship between early word components and later comprehension components.

Combined, these patterns suggest that JC3 marks the point where the left TPJ cluster shifts from connectivity with the language network to connectivity with the DMN, potentially reflecting a “bottom-up” spreading activation linking the retrieved word to the larger context of the situation model (Humphreys et al., 2021; Smallwood & Schooler, 2015; Kveraga et al., 2007). Consistent with this interpretation, isolated Word reading did not show a robust relationship with this component. This suggests that 500 ms after LC stimuli, there is a marked change between contextualized and non-contextualized language processes. The theory that JC3 may flexibly support a transition from local (language) and global (inferential) processes may help resolve competing findings regarding the Late Positive Component, specifically the debate about whether the early P600 reflects domain-specific processing (e.g., Friederici et al. (1996)) or domain-general processes (Kolk & Chwilla, 2007; Kuperberg, 2007).

The two final components, JC4 and JC5, fell within the canonical P600 window. JC4 localized to a semantic control network, integrating hubs from JC2 and JC3 with activation in the left IFG (Jefferies, 2013). The repeated involvement of the left TPJ across JC2-JC4 underscores its role as a dynamic hub facilitating within- and cross-network coupling during distinct temporal windows of LC (Seghier, 2013). Furthermore, the text feature analysis suggested that this differential coupling promotes different cognitive processes, with the JC4 network being related to more difficult visualization of language (consistent with prior P600 work; Brouwer et al. (2012); Petten & Luka (2012); Molinaro et al. (2011)). The spatial, temporal and linguistic profile of this component is consistent with literature on binding information during complex language integration. The last component, JC5, localized to core DMN and FPN nodes in Passages and was dependent on greater visualization ease of the language, and is consistent with a higher-order cross-network connectivity previously found to support processes related to mental model construction in language (Aboud et al., 2023; Fedorenko, 2014). This component was distinguished by medial and lateral prefrontal activations; the medial prefrontal cortex is associated with constructing and integrating abstract representations (Blumenfeld & Ranganath, 2019; Gilboa et al., 2004) and lateral prefrontal regions linked to goal-directed executive processes (Duncan, 2010; Miller & Cohen, 2001). JC5 may reflect the culmination of earlier processes or serve as a predictive mechanism preparing for upcoming text units, underscoring its role in maintaining and updating coherent mental representations.

### LC Efficiency and Individual Differences

We next questioned which parts of this cascade were most strongly related to efficient LC processing. We found that high LC performance is marked by (1) less reliance on the semantic control network (left IFG, left MTG, left AG), (2) greater inferential posterior DMN relative to the semantic control network, and (3) more mediation of early and late signals by the the posterior DMN network. Together, we find that LC efficiency is marked by the interplay between spreading activations in the default mode and engagement of top-down control. Our findings are consistent with studies on expertise: cognitive efficiency is often marked by more parsimonious reliance on explicit cognitive control mechanisms and increased spreading activation (Daselaar et al., 2015; Neubauer & Fink, 2009). This reduced need for overt cognitive control in high LC individuals could reflect the development of highly tuned/specialized LC pathways, where comprehension unfolds in a more automatic manner. Conversely, less efficient but still successful LC (as captured by the current cohort), may require increased reliance on top-down semantic control mechanisms to drive LC processes when inference is not enough to guide understanding. This interpretation is consistent with prominent LC theories (The Landscape Model, Yeari & van den Broek (2011)), which suggest that strong comprehension is characterized by the real-time nuanced balance between inferential processes connecting language content and background knowledge, and top-down assessment processes to help resolve ambiguities and maintain coherence (Perfetti, 2007). Future work should examine how these patterns differ in adults with low LC ability.

### Limitations and Future Directions

While our stimuli varied along key text dimensions related to LC, syntactic complexity was controlled. Future studies should expand this multimodal characterization to examine the impact of syntactic variability and anomalies. Additionally, single-word presentation minimized artifacts and optimized signal clarity but likely introduced additional working memory demands compared to naturalistic text presentations. Examining these dynamics under more naturalistic conditions will be critical for validating and extending these findings. Finally, while we linked spatiotemporal signals to cognitive demands through linguistic features, further research is needed to disentangle each component’s sensitivity to specific language dimensions, such as social versus scientific text differences.

The current study presents a methodological and conceptual framework for examining the rapid neural dynamics of naturalistic LC. By leveraging fused brain imaging analytics, our findings provide evidence for a specific progression of brain networks underlying LC and establish a platform for future investigations into brain network dynamics across clinical and cognitive domains. By situating LC within broader principles of distributed cognition and dynamic network integration, we aim to advance our understanding of complex cognitive processes and their neural underpinnings.

## Appendix A

Illustration of joint ICA

**Figure A1.**
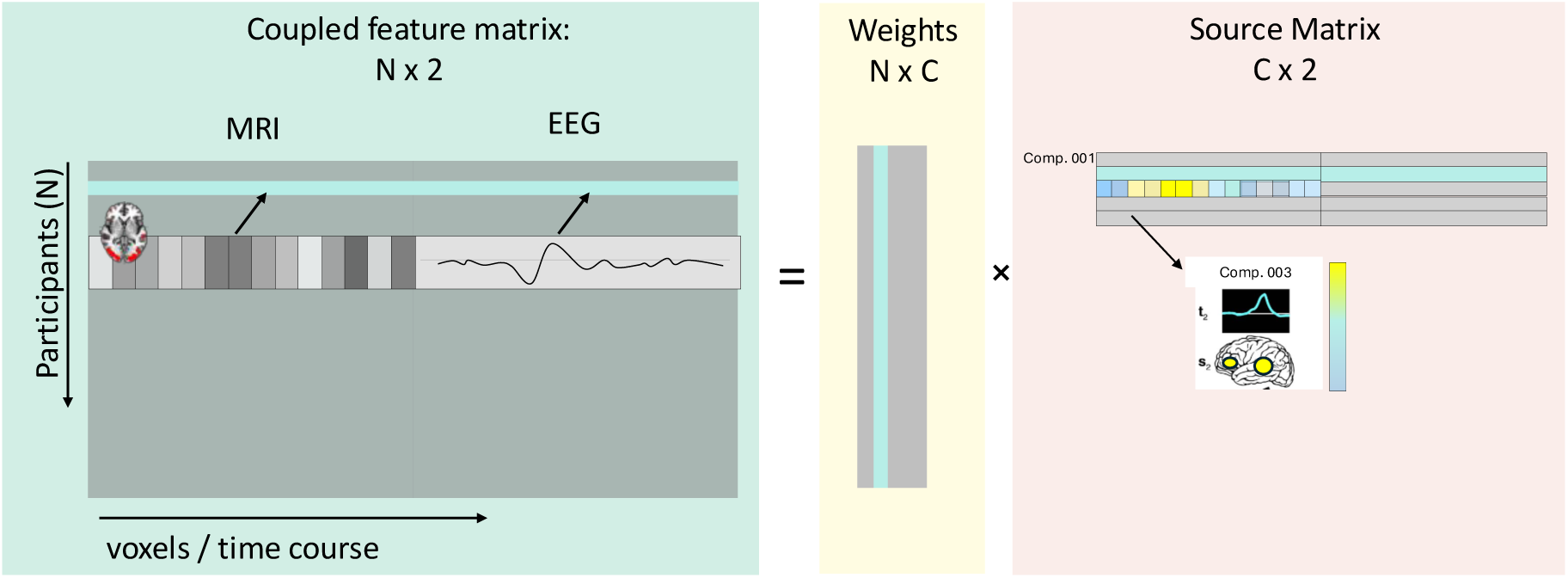
Joint Independent Component Analysis: A simplified schematic representation of joint Independent Component Analysis (ICA). This approach integrates EEG and fMRI data from multiple participants into a unified matrix for simultaneous analysis. The joint ICA procedure captures signal variations across both subjects and imaging modalities, enabling the identification of distinct neural patterns (i.e., independent components). These components represent spatial information derived from fMRI and temporal patterns obtained from EEG, providing a comprehensive view of brain activity. The resulting analysis leverages the high spatial resolution of fMRI and the excellent temporal resolution of EEG, offering insights into both where and when neural processes occur during reading.

## Appendix B

**Table B1.** Spatial Activations for Passage Joint Components.

| Area | BA | Cluster size | T-value | MNI coordinates (x, y, z) |
| --- | --- | --- | --- | --- |
| Passage Joint Comp. 1: N400a |  |  |  |  |
| L Mid Occipital | 19 | 2445 | 5.8 | (-19.7, -99, 7.1) |
| R Precentral | 4 | 551 | 4.6 | (50.3, -2.4, 47.0) |
| L Fusiform | 37 | 265 | 4.1 | (-41.1, -55.5, -15.0) |
| R Inf Frontal Tri | 45 | 113 | 3.7 | (47.4, 38.2, 10.8) |
| R Mid Occipital | 19 | 102 | 4.3 | (31.2, -91.6, 13.0) |
| L Mid Occipital | 19 | 82 | 3.7 | (-51.4, -72.4, -1.0) |
| L Mid Frontal | 46 | 70 | 3.3 | (-41.1, 46.3, 16.0) |
| L Postcentral | 3 | 66 | 3.6 | (-57.3, -9.0, 41.8) |
| L Precentral | 4 | 56 | 4.2 | (-44.8, -4.6, 55.1) |
| R Sup Temporal | 41 | 39 | 3.4 | (67.3, -28.9, 18.9) |
| R Cerebellum Crus I | - | 32 | 3.5 | (12.7, -87.9, -23.8) |
| L Cuneus | 17 | 32 | 4.0 | (-11.6, -93.1, 33.0) |
| Passage Joint Comp. 2: N400b |  |  |  |  |
| R Calcarine | 17 | 927 | 7.2 | (16.4, -94.5, -1.0) |
| R Mid Occipital | 19 | 629 | 4.9 | (42.2, -82.0, 3.5) |
| R Sup Occipital | 19 | 524 | 5.9 | (23.1, -92.3, 18.2) |
| L Mid Occipital | 19 | 447 | 5.5 | (-38.2, -84.2, -0.2) |
| L Mid Temporal | 21 | 110 | 4.1 | (-58.8, -57.7, 17.5) |
| Passage Joint Comp. 3: P600a |  |  |  |  |
| L Mid Occipital | 19 | 1487 | 5.7 | (-43.3, -82.0, -3.9) |
| R Mid Temporal | 39 | 1020 | 4.3 | (45.2, -75.4, 5.7) |
| R Sup Occipital | 19 | 485 | 5.5 | (21.6, -93.8, 17.5) |
| L Calcarine | 17 | 481 | 5.0 | (-17.5, -101.2, -1.7) |
| R Mid Temporal | 21 | 296 | 4.2 | (45.9, -40.7, 10.1) |
| R Sup Temporal | 22 | 280 | 4.0 | (63.6, -43.7, 17.5) |
| L Postcentral | 3 | 228 | 4.0 | (-54.4, -9.7, 44.0) |
| R Mid Temporal | 21 | 225 | 3.9 | (58.5, -57.7, 14.5) |
| R Angular | 39 | 221 | 3.5 | (40.0, -62.1, 19.7) |
| L Postcentral | 3 | 169 | 3.4 | (-55.1, -6.0, 24.8) |
| R Sup Frontal | 10 | 145 | 3.2 | (30.4, 57.4, 8.6) |
| L Mid Occipital | 19 | 139 | 3.7 | (-31.5, -94.5, 4.9) |
| R Mid Occipital | 19 | 135 | 3.3 | (31.2, -73.2, 27.8) |
| L Mid Temporal | 22 | 103 | 4.3 | (-61.8, -54.7, 16.0) |
| R Inf Temporal | 37 | 94 | 3.8 | (53.3, 30.8, -31.9) |
| R Mid Frontal | 9 | 93 | 3.3 | (28.2, 52.2, -0.2) |
| R Lingual | 18 | 87 | 5.3 | (17.9, -101.9, -6.9) |
| R Cerebellum 6 | - | 74 | 3.5 | (37.8, -59.1, -19.4) |
| R Mid Occipital | 19 | 73 | 3.2 | (34.1, -83.5, 30.0) |
| R Mid Frontal | 9 | 63 | 3.2 | (31.9, 30.8, 36.6) |
| R Precuneus | 7 | 55 | 3.3 | (11.3, -56.9, 26.3) |
| L Mid Occipital | 19 | 43 | 3.6 | (-28.6, -87.9, 30.7) |
| R Inf Frontal Orb | 11 | 37 | 3.3 | (28.2, 30.1, -16.5) |
| Passage Joint Comp. 4: P600b |  |  |  |  |
| L Calcarine | 18 | 1863 | - | (-8.7, -90.1, -8.3) |
| R Cerebellum Crus I | - | 1700 | 4.2 | (18.6, -78.3, -29.7) |
| L Superior Parietal | 7 | 699 | 5.4 | (-18.2, -76.8, 47.7) |
| R Lingual | 18 | 648 | 4.5 | (14.2, -88.6, -1.7) |
| L Calcarine | 17 | 320 | 3.5 | (-15.3, -72.4, 7.9) |
| L Cerebellum VI | - | 180 | 3.8 | (-35.9, -67.3, -18.7) |
| L Middle Temporal | 21 | 180 | 3.9 | (-49.2, -51.8, 16.7) |
| L Middle Temporal | 21 | 145 | 3.6 | (-58.8, -28.9, -0.2) |
| Vermis VII | - | 117 | 3.8 | (5.4, -77.6, -23.8) |
| L Precentral | 6 | 85 | 3.8 | (-44.1, 9.4, 49.9) |
| R Middle Occipital | 18 | 45 | 3.2 | (30.4, -91.6, 9.4) |
| R Superior Parietal | 7 | 37 | 3.4 | (13.5, -62.8, 58.8) |
| L Inferior Occipital | 18 | 34 | 3.4 | (-27.1, -83.5, -3.2) |
| L Lingual | 18 | 33 | 3.7 | (-24.9, -92.3, -15.0) |
| Passage Joint Comp. 5: P600c |  |  |  |  |
| L Postcentral | 3 | 928 | 3.6 | (-55.9, -6.0, 26.3) |
| L Angular | 39 | 669 | 3.5 | (-36.7, -58.4, 33.0) |
| R Superior Medial Frontal | 10 | 652 | 3.2 | (10.5, 61.8, 11.6) |
| R Middle Frontal | 46 | 526 | 3.3 | (39.3, 47.8, 15.3) |
| R Middle Frontal | 46 | 324 | 3.2 | (28.2, 37.5, 41.1) |
| R Anterior Cingulate | 32 | 191 | 2.9 | (6.1, 45.6, 11.6) |
| R Superior Parietal | 7 | 130 | 3.1 | (20.1, -45.1, 69.1) |
| R Superior Frontal | 8 | 126 | 3.0 | (18.6, 27.9, 54.3) |
| R Precuneus | 31 | 89 | 2.8 | (9.0, -56.9, 22.6) |
| L Middle Frontal | 10 | 60 | 2.8 | (-33.0, 56.6, 7.1) |
| R Middle Frontal | 9 | 59 | 2.9 | (39.3, 36.0, 35.2) |

## Appendix C

**Table C1.** Spatial Activations for Words Joint Components.

| Area | BA | Cluster size | T-value | MNI coordinates (x, y, z) |
| --- | --- | --- | --- | --- |
| Words Joint Comp. 1: N400a |  |  |  |  |
| L Middle Occipital | 19 | 2296 | 7.1 | (-30.8, -90.9, 4.2) |
| R Middle Occipital | 19 | 2127 | 6.1 | (29.0, -90.9, 7.9) |
| R Superior Occipital | 19 | 73 | 4.9 | (20.8, -93.8, 24.8) |
| L Middle Temporal | 21 | 223 | 4.5 | (-62.5, -31.9, 6.4) |
| L Middle Occipital | 17 | 123 | 4.5 | (-8.7, -102.7, 0.5) |
| Words Joint Comp. 2: N400b |  |  |  |  |
| L Middle Temporal | 21 | 1620 | 5.3 | (-53.6, -40.7, 6.4) |
| L Inferior Frontal Operculum | 44 | 1151 | 5.2 | (-41.8, 10.9, 24.8) |
| L Middle Occipital | 19 | 401 | 3.5 | (-41.8, -75.4, 2.7) |
| L Postcentral | 3 | 357 | 4.3 | (-61.0, -3.8, 21.9) |
| R Calcarine | 17 | 247 | 3.7 | (15.7, -75.4, 9.4) |
| L Superior Temporal Pole | 38 | 203 | 4.1 | (-50.0, 15.3, -11.3) |
| L Superior Medial Frontal | 9 | 203 | 3.9 | (-1.3, 46.3, 49.9) |
| L Inferior Frontal Triangularis | 45 | 153 | 3.8 | (-53.6, 30.1, 2.0) |
| R Rolandic Operculum | 43 | 152 | 3.9 | (62.9, -2.4, 14.5) |
| L Fusiform | 37 | 147 | 3.3 | (-45.5, -56.2, -16.5) |
| L Inferior Parietal | 40 | 114 | 3.6 | (-30.0, -48.1, 45.5) |
| R Middle Temporal | 21 | 95 | 3.3 | (43.0, -62.8, 7.9) |
| L Superior Temporal | 22 | 80 | 3.8 | (-58.8, 3.5, -6.9) |
| L Middle Frontal | 46 | 52 | 3.1 | (-32.3, 48.5, 9.4) |
| L Superior Occipital | 19 | 52 | 3.1 | (-21.2, -79.1, 23.4) |
| L Superior Medial Frontal | 9 | 49 | 3.0 | (-2.8, 40.4, 35.2) |
| R Middle Temporal Pole | 38 | 43 | 3.7 | (51.1, 17.6, -29.0) |
| L Middle Cingulate | 31 | 40 | 3.1 | (-14.6, -49.6, 33.7) |
| L Middle Temporal | 21 | 39 | 2.8 | (-62.5, -48.1, 10.8) |
| L Middle Frontal | 6 | 32 | 2.9 | (-40.4, 2.1, 54.3) |
| Words Joint Comp. 3: P600a |  |  |  |  |
| L Middle Temporal | 21 | 1248 | 4.4 | (-58.1, -30.4, 1.2) |
| L Cuneus | 18 | 1207 | 3.9 | (-2.8, -81.3, 35.2) |
| L Middle Occipital | 19 | 866 | 4.8 | (-39.6, -86.4, -4.7) |
| L Precentral | 6 | 402 | 3.7 | (-47.0, 3.5, 47.0) |
| R Middle Occipital | 19 | 374 | 4.0 | (44.4, -82.0, 3.5) |
| R Cuneus | 18 | 288 | 3.7 | (16.4, -70.2, 38.9) |
| R Superior Occipital | 19 | 231 | 3.9 | (23.1, -93.1, 21.9) |
| L Middle Temporal | 21 | 148 | 3.6 | (-48.5, -48.8, 6.4) |
| L Inferior Frontal Triangularis | 45 | 142 | 3.3 | (-39.6, 38.2, 8.6) |
| L Inferior Frontal Operculum | 44 | 126 | 3.6 | (-33.7, 5.8, 33.0) |
| L Superior Temporal | 22 | 97 | 3.8 | (-64.7, -43.7, 16.0) |
| L Superior Medial Frontal | 9 | 91 | 3.6 | (-7.2, 59.6, 33.0) |
| L Superior Occipital | 19 | 79 | 3.3 | (-24.9, -73.2, 26.3) |
| R Middle Occipital | 19 | 71 | 3.2 | (39.3, -67.3, 6.4) |
| R Cerebellar Crus II | - | 47 | 3.3 | (2.4, -84.2, -29.7) |
| L Inferior Frontal Operculum | 44 | 40 | 2.8 | (-43.3, 12.4, 27.8) |
| L Inferior Frontal Triangularis | 45 | 40 | 2.8 | (-41.1, 17.6, 23.4) |
| Cerebellar Tonsil | - | 39 | 3.0 | (20.8, -49.6, -36.4) |
| L Supplementary Motor Area | 6 | 39 | 3.0 | (-1.3, 17.6, 57.3) |
| L Cerebellum VI | - | 38 | 3.0 | (-7.9, -69.5, -6.9) |
| L Supplementary Motor Area | 6 | 32 | 3.1 | (-2.0, 9.4, 65.4) |
| Words Joint Comp. 4: P600b |  |  |  |  |
| L Lingual | 18 | 4531 | 4.5 | (-7.9, -85.0, -10.6) |
| L Superior Parietal | 7 | 505 | 4.2 | (-19.0, -68.7, 55.1) |
| L Superior Occipital | 19 | 401 | 5.0 | (-16.0, -79.8, 45.5) |
| R Middle Occipital | 19 | 284 | 4.0 | (39.3, -82.0, 15.3) |
| R Calcarine | 17 | 249 | 3.6 | (17.9, -73.2, 7.9) |
| R Cerebellar Lobule VIIb | - | 245 | 3.3 | (37.1, -63.6, -48.2) |
| L Calcarine | 17 | 244 | 3.6 | (-4.2, -75.4, 10.1) |
| L Calcarine | 17 | 165 | 3.3 | (-20.5, -64.3, 4.9) |
| L Middle Occipital | 19 | 128 | 3.4 | (-36.7, -83.5, 11.6) |
| R Precentral | 6 | 127 | 3.5 | (40.0, 2.8, 41.8) |
| L Superior Medial Frontal | 10 | 127 | 3.4 | (-8.7, 59.6, 7.9) |
| R Fusiform | 19 | 122 | 3.2 | (20.1, -74.6, -13.5) |
| L Inferior Occipital | 19 | 103 | 3.1 | (-44.1, -68.0, -3.9) |
| R Cerebellum VI | - | 88 | 3.0 | (31.9, -69.5, -22.4) |
| L Middle Occipital | 18 | 66 | 3.3 | (-28.6, -90.1, 18.2) |
| L Middle Occipital | 19 | 59 | 3.2 | (-23.4, -86.4, 5.7) |
| Words Joint Comp. 5: P600c |  |  |  |  |
| R Middle Frontal | 9 | 1444 | 4.1 | (34.9, 33.8, 38.1) |
| R Superior Frontal | 10 | 354 | 3.1 | (26.7, 55.2, 0.5) |
| R Middle Frontal | 46 | 339 | 3.5 | (34.9, 54.4, 18.2) |
| L Postcentral | 3 | 239 | 3.7 | (-61.8, -4.6, 18.2) |
| L Postcentral | 3 | 178 | 3.5 | (-64.7, -15.6, 25.6) |
| R Angular | 39 | 134 | 3.2 | (46.7, -70.2, 35.2) |
| L Inferior Parietal | 40 | 72 | 2.9 | (-58.8, -42.2, 40.3) |

